# An adjuvant for in vivo CAR-T therapy potentiation against solid tumors

**DOI:** 10.64898/2026.10.30.763005

**Authors:** Yu Sun, Yishan Ma, Tong Guan, Yumeng Yan, Yali Ming, Xiaonan Huang, Yujie Luo, Jingsheng Cai, Guanghui Ma, Fuping You, Mantang Qiu, Jingyi Chen, Yufei Xia

## Abstract

*In vivo* CAR-T therapy offers rapid and controllable T-cell engineering but has yet to achieve durable efficacy against solid tumors. Rather than focusing solely on CAR-T cells, we harness the intact immune system via an *in vivo* CAR-T adjuvant (iCTA). Composed of aluminum– manganese hydrogel, iCTA can form a spatially confined antigenic and immunostimulatory niche *in-situ*. By coupling controlled antigen exposure with STING-type I interferon signaling, iCTA not only recruited and reactivated CAR-T cells, but engage the CAR-negative immunocytes with enhanced dendritic-cell antigen presentation and mobilized endogenous antigen-specific T-cell responses. Combined with mRNA-LNP-mediated *in vivo* CAR-T generation, a single administration of iCTA increased CAR⁺ T-cell abundance in draining lymph nodes and blood by 2-fold. It also increased T and NK cell infiltration, established an IFNγ-enriched tumor microenvironment, and reduced exhaustion-associated T-cell states. iCTA improved tumor regression and survival, establishing an adjuvant strategy to enhance *in vivo* CAR-T therapy.

## INTRODUCTION

*In vivo* CAR-T therapy can directly program endogenous T cells to express CARs *in situ*,^1–3^ potentiating the elimination of pathogenic B cells or tumor cells,^4,5^ while bypassing the complex manufacturing required for conventional *ex vivo* cell therapy.^6^ Transient, non-integrating gene delivery, such as mRNA-encapsulated lipid nanoparticles (mRNA-LNPs),^3,7^ may further improve controllability by avoiding the permanent genomic modification associated with viral transduction. However, transient CAR expression and endogenous T cell expansion may constrain the magnitude and persistence of *in vivo*-generated CAR-T cells, particularly against established tumors. Moreover, CAR-positive T cells may receive insufficient or poorly coordinated antigenic and costimulatory support in peripheral tissues, limiting sustained expansion and effector function.^8,9^ Within the tumor, persistent antigen exposure and an immunosuppressive tumor microenvironment (TME) further accelerate functional exhaustion, hampering the antitumor effects.^10–12^

Efforts to improve *in vivo* CAR-T therapy have largely focused on enhancing T-cell targeting, transfection, and direct stimulation via mRNA and LNP engineering.^13–15^ However, intensifying or prolonging T-cell stimulation risks premature differentiation and exhaustion.^16,17^ Without preparative lymphodepletion,^17–19^ *in vivo* CAR-T engineering can preserve an intact endogenous immune system, creating an additional opportunity to engage immune populations beyond the CAR-positive fraction. Rather than activating this immune network as a parallel therapy, we hypothesized that it could be harnessed to boost *in vivo* CAR-T cell responses and broaden antitumor activity beyond the engineered T cell populations. Under these circumstances, dendritic cells (DCs), helper T cells, and NK cells, could provide complementary antigen presentation, immune activation, and TME remodeling.

Here, we developed an *in vivo* CAR-T adjuvant (iCTA) based on an aluminum-manganese composite hydrogel. By coordinating controlled antigen exposure with cGAS-STING signaling, iCTA established a local niche *in situ*, that recruited and reactivated CAR-positive T cells, while propagating immune activation to the endogenous immune system. Following administration, iCTA increased CAR-positive T cell abundance in draining lymph nodes and blood. Moreover, iCTA improved antigen-presentation of DCs and stimulated the antigen-specific T cell responses, while reshaping TME into a high-IFNγ state with more T and NK cell engagement (**Figure 1A**). In DLL3-expressing small-cell lung cancer models, combining iCTA with *in vivo* CAR-T therapy improved tumor control and survival. This adjuvant strategy may extend *in vivo* CAR-T therapy beyond engineering CAR-T cells alone, towards coordinated modulation of the immune system.

**Figure 1.**
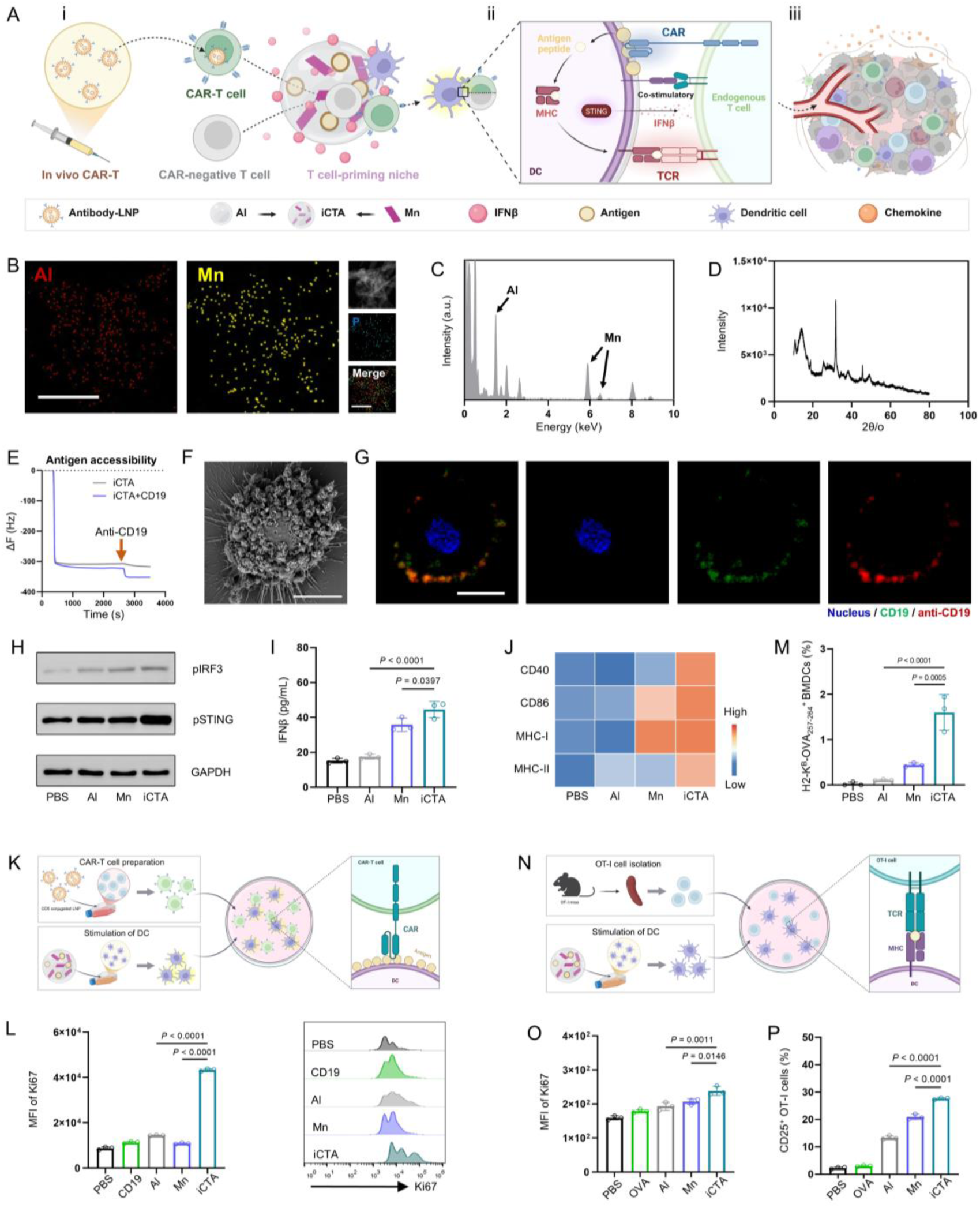
Preparation of iCTA and evaluation of the activation of endogenous T cells *in vitro*. (A) Proposed mechanism of the *in vivo* CAR-T adjuvant (iCTA). (i) Intravenous delivery of antibody-conjugated LNPs generates CAR-T cells *in vivo*. Subsequent subcutaneous iCTA administration recruits endogenous CAR-T and CAR-negative T cells to form a local T cell-priming niche. (ii) Within this niche, alum (Al) and manganese (Mn) cooperatively activate cGAS-STING signaling of antigen-presenting cells (APCs). Dendritic cells (DCs) present antigen to CAR-T cells, while iCTA also enhances DC cross-presentation to prime endogenous T cells. (iii) iCTA reshapes the tumor microenvironment (TME) and promotes further T-cell infiltration. (B) Representative elemental mapping images of iCTA. Scale bar, 500 nm. (C) EDS elemental analysis of iCTA. Arrows indicate the characteristic peaks of Mn. (D) XRD analysis of iCTA. (E) QCM analysis of CD19 accessibility. iCTA or CD19-loaded iCTA was immobilized on gold sensor chips. Anti-CD19 antibody was introduced at 44 min. (F) SEM images of iCTA adsorbed on DC2.4 cell surfaces after 6 h co-incubation. Scale bar, 5 μm. (G) Confocal images of CD19 antigen (green) and anti-CD19 antibody (red) adsorption on hBMDC membranes after 6 h. Scale bar, 5 μm. (H) Western blot analysis of pIRF3, pSTING, and GAPDH in BMDCs 24 h after adjuvant treatment. (I) ELISA quantification of IFNβ concentrations in the supernatant of BMDCs 24 h after stimulation with iCTA (n = 3). (J) Flow cytometric heatmap of CD40, CD86, MHC-I, and MHC-II expression on BMDCs 24 h after stimulation (n = 3). (K) Schematic illustration of *in vitro* CAR-T activation. hBMDCs were stimulated with indicated adjuvants and co-cultured with CD19 CAR-T cells for 48 h. (L) CAR-T proliferation by flow cytometry. Ki67 MFI and representative histograms are shown (n= 3). (M) Surface H-2K^B^–SIINFEKL complexes on mBMDCs after 24 h stimulation with OVA plus indicated adjuvants (n = 3). (N) Schematic illustration of *in vitro* antigen-specific activation. Adjuvant-stimulated mBMDCs were co-cultured with OT-I cells for 48 h. (O, P) OT-I activation by flow cytometry. (O) Ki67 MFI and (P) CD25 expression level (n = 3). Data in (I), (L), (M), (O), and (P) are presented as the mean ± SD. Statistical significance was calculated via one-way ANOVA with a Tukey post-hoc test.

## RESULTS

### Engineering iCTA for enhanced CAR-T cell and antigen-specific T cell responses

To construct an immunostimulatory niche capable of supporting both CAR recognition and broader T-cell priming, we sought to integrate sustained antigen availability with localized innate immune activation within a single material scaffold.^20^ Aluminum hydroxide (Al), a clinically established adjuvant, was repurposed as a locally retained, gel-like scaffold^21–23^ that enables controlled antigen exposure and forms an antigen depot for immune cell recruitment. In addition, manganese (Mn) was incorporated to provide cGAS-STING-dependent immune stimulation.^24,25^ To spatially couple these functions, aluminum and manganese chlorides were co-precipitated under pH-stable conditions to generate a co-condensed Al-Mn composite, in which antigen retention and innate immune activation were physically integrated.

We next screened a series of Al to Mn ratios to balance immune activation against excessive inflammatory signaling. A 1:1 mass ratio elicited the highest interferon β (IFNβ) level while maintaining minimal TNFα secretion and was therefore identified as the optimal iCTA ratio (**Figure S1A**). The element mapping images and energy dispersive spectroscopy (EDS) analysis showed that iCTA consisted of Al and Mn (**Figures 1B** and **1C**) with an average particle size of approximately 1.4 μm and a positively charged surface (**Figures S1B** and **S1C**). X-ray diffraction (XRD) structural characterization and transmission electron microscopy (TEM) revealed that iCTA exhibited a porous, mesh-like amorphous structure, supporting its potential to form an immune-cell-recruiting niche after injection (**Figures 1D** and **S1D**). X-ray photoelectron spectroscopy (XPS) further showed that Mn retained its valence state following formulation (**Figure S1E**), while iCTA induced no observable cytotoxicity in bone marrow-derived dendritic cells (BMDCs, **Figure S1F**). The above results indicated that iCTA preserved the depot-forming and antigen-loading features of alum while incorporating additional innate immunostimulatory effects.

Given that CAR activation requires extracellular accessibility of its cognate antigen,^26,27^ we next asked whether antigen retained by iCTA remained available for direct CAR recognition. Quartz crystal microbalance (QCM) measurements demonstrated iCTA-immobilized CD19 retained antibody-binding capacity (**Figure 1E**), suggesting antigen accessibility for direct CAR interaction. We then assessed whether this accessible antigen could be presented at the cell surface. Scanning electron microscopy (SEM) revealed close association of iCTA with the DC plasma membrane (**Figure 1F**), while confocal laser scanning microscopy (CLSM) further showed the membrane-localized iCTA and associated antigen with BMDCs (**Figures S1G and S1H**). Consistently, surface immunostaining of human BMDCs incubated with CD19-loaded iCTA demonstrated that scaffold-bound CD19 remained accessible to anti-CD19 antibody at the cell surface (**Figure 1G**). Together, these findings showed that iCTA preserves antigen accessibility both on the scaffold and after membrane association, supporting direct CAR engagement and potential antigen recognition of the APCs.

Immunoblotting and enzyme-linked immunosorbent assay (ELISA) demonstrated robust cGAS-STING activation in iCTA-treated BMDCs (**Figures 1H** and **1I**). iCTA also increased the expression of the co-stimulatory molecules CD40 and CD86 and enhanced surface MHC-I and MHC-II expression (**Figure 1J**), indicating coordinated DC maturation and increased antigen presentation. Thus, iCTA integrates the complementary functions of accessible antigen display for CAR recognition and APC activation for advanced T-cell priming.

We next tested whether this dual functionality translated into T-cell activation. hBMDCs pulsed by CD19-loaded iCTA (iCTA^CD^^19^) showed increased Ki67 expression in co-cultured CD19-CAR-T cells (**Figures 1K** and **1L**), consistent with enhanced CAR-T cell proliferation. Thus, iCTA couples APC activation and accessible CAR antigen to support CAR-T cell expansion. In addition to CAR-positive cells, we also evaluated antigen-specific activation of CAR-negative T cells using OVA as a model antigen. As shown in **Figures 1M** and **S1I**, iCTA^OVA^ increased presentation of the H-2K^B^–SIINFEKL complex on DCs and expanded antigen-specific CD8^+^ T cells detected by tetramer staining. Consistently, OT-I T cells exhibited enhanced antigen-specific priming and proliferation (**Figures 1N-1P** and **S1J**). These findings indicated that iCTA harbored the potential to induce direct CAR-T reactivation, and APC-mediated, TCR-dependent T-cell priming.

### iCTA establishes a T-cell-priming niche *in vivo*

We next evaluate whether iCTA could establish a spatially organized T-cell-priming niche *in vivo*. Bioluminescence imaging showed that iCTA retained antigen at the injection site for more than seven days (**Figure 2A**), establishing a durable local antigen depot. Then, subcutaneous iCTA nodules were excised for histological and ultrastructural analyses (**Figure 2B**). Hematoxylin and eosin (H&E) histological examination (**Figure 2C**) and SEM images (**Figure 2D**) demonstrated dense cellular infiltration within the nodules. After identifying 24 h as the optimal time point for T cell recruitment by flow cytometry (**Figure S2A**), we performed single-cell RNA sequencing (scRNA-seq) on CD45⁺ cells isolated from the injection site at this time point. Unsupervised clustering identified seven major immune populations including T cells, macrophage/monocytes (macro/monos), DCs, and neutrophils (**Figures 2E, 2F** and **S2B**), among which the recruitment of T cells was associated with chemokine programs expressed by innate immune cells (**Figure S2C**).

**Figure 2.**
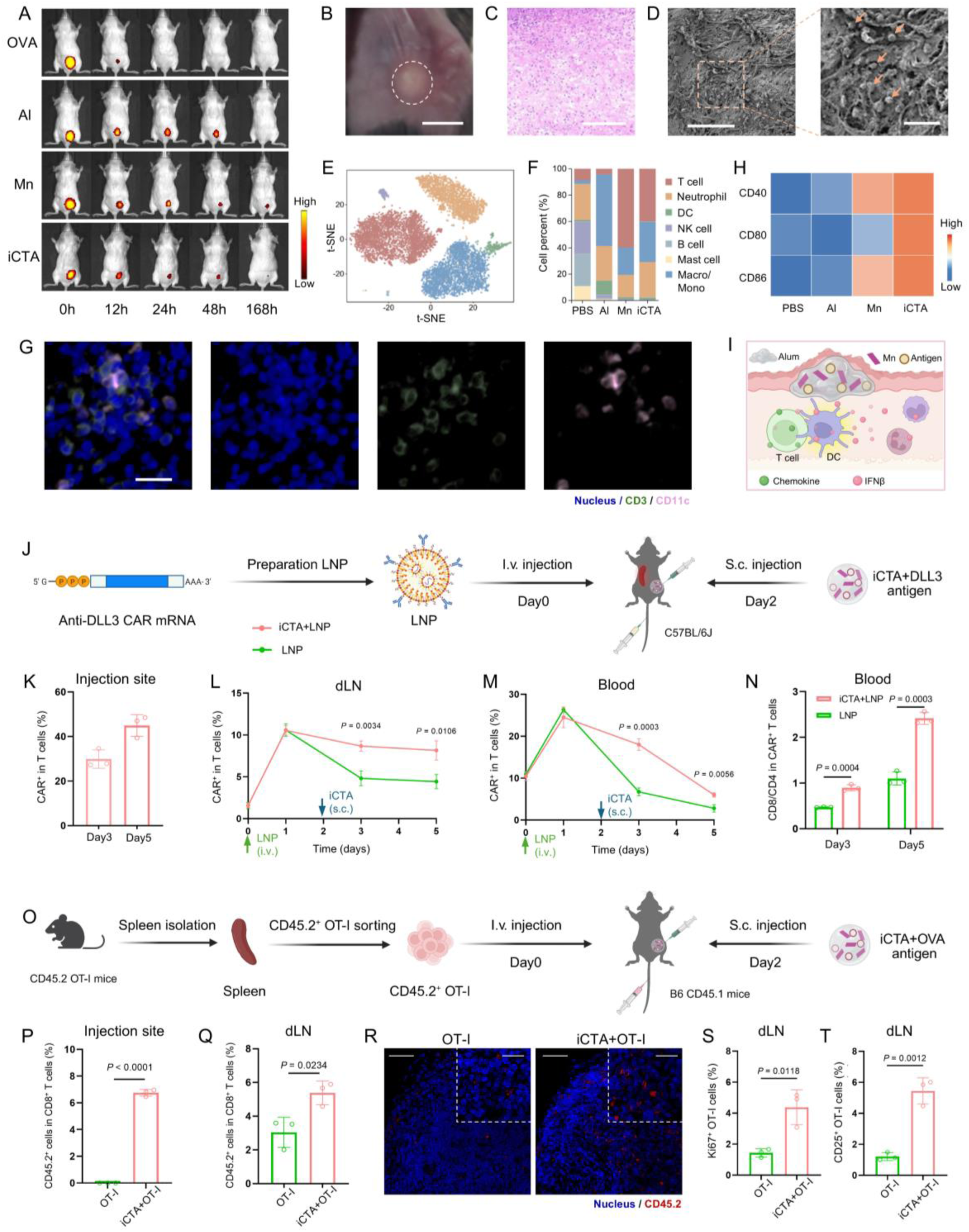
Evaluation of the establishment of T cell-priming niche *in vivo*. (A) Retention of Cy5-labeled antigen at the injection site after subcutaneous administration of indicated antigen-adsorbed adjuvants, monitored from 0 to 168 h. (B) Image of the injection site 24 h after iCTA administration. Scale bar, 2 mm. (C) Representative histology of injection-site nodules on day 1 after iCTA injection. Scale bar, 200 μm. (D) SEM image of nodules excised from injection sites 24 h after indicated treatments. Left scale bar, 80 μm; right scale bar, 20 μm. (E, F) Mice received subcutaneous injection of antigen-adsorbed adjuvants. Tissues were collected on day 1 for scRNA-seq. (E) t-SNE projection and (F) cell-type proportions. (G) Representative immunofluorescence of CD3^+^ T cells (green) and CD11c^+^ DCs (pink) at iCTA injection sites. Scale bar, 50 μm. (H) Flow-cytometric heatmap of CD40, CD80, and CD86 on DCs in injection-site tissue at 24 h (n= 3). (I) Schematic of iCTA-driven *in situ* T cell-priming niche formation. (J) Experimental design for *in vivo* CAR-T expansion with iCTA^DLL3^. LNPs were given intravenously on day 0, iCTA^DLL3^ subcutaneously on day 2, and tissues collected at indicated time points for flow cytometry. (K-M) Longitudinal frequencies of CAR^+^ T cells at injection site (K), draining lymph node (dLNs, L), and blood (M), comparing iCTA + LNP versus LNP alone (n = 3). (N) The proportions of CD8⁺ and CD4⁺ T cells among CAR⁺ T cells in peripheral blood at days 3 and 5 (n = 3). (O) Schematic illustration of the experimental design used to OT-I trafficking after iCTA^OVA^. CD45.2^+^ OT-I cells were transferred on day 0; iCTA^OVA^ was injected on day 2; tissues were harvested on day 5. (P) CD45.2^+^ OT-I recruitment to iCTA injection sites by flow cytometry (n = 3). (Q) CD45.2^+^ OT-I accumulation in dLNs. (n = 3). (R) Immunofluorescence of CD45.2^+^ OT-I cells (red) in dLNs. Left scale bar: left, 50 μm; right scale bar, 20 μm. (S, T) Functional status of CD45.2^+^ OT-I cells in dLNs: (S) Ki67 proliferation and (T) CD25 activation (n = 3). Data in (K-N), (P), (Q), (S) and (T) are presented as the mean ± SD. Statistical significance was calculated via two-tailed Student’s *t* test (L, M, N, P, Q, S, and T).

We next assessed interactions between DCs and T cells within the local niche.^28^ CLSM imaging revealed co-localization of DCs and T cells at the iCTA injection site (**Figure 2G**). Additionally, ligand-receptor interaction analysis on the scRNA-seq dataset showed that the number of DC-CD8⁺ T cell interactions was markedly higher in the iCTA group than in the Al or Mn groups, regardless of which cell type served as ligand or receptor (**Figure S2D**). Further classification revealed prominent antigen-presentation axes (MHC-II–CD4 and MHC-I–CD8) and canonical CD80/CD86–CD28 co-stimulatory signaling (**Figure S2E** and **S2F**). Moreover, ELISA revealed elevated local IFNβ secretion following iCTA injection in the niche (**Figure S2G**), suggesting an immunostimulatory microenvironment.

Functionally, DCs within the iCTA niche displayed marked upregulation of CD40, CD80, and CD86 (**Figure 2H**), further providing co-stimulatory signal for T cell activation. The DCs also exhibited enhanced antigen processing and presentation, as shown by increased surface H-2K^B^–SIINFEKL complexes (**Figure S2H**). iCTA triggered an elevated expansion of OVA-tetramer CD8^+^ T cells in draining lymph nodes (dLNs, **Figure S2I**), indicating increased generation of antigen-specific T cells. Moreover, we found elevated expression of MHC-II, corroborated by bulk RNA-seq of sorted CD11c⁺ DCs (**Figure S2J**).

Together, these results demonstrate that iCTA establishes a spatially organized T-cell-priming niche that combines prolonged antigen retention, immunocyte recruitment, DC activation and antigen presentation to promote local T-cell activation and systemic antigen-specific cellular immunity (**Figure 2I**).

### Recruiting *in vivo*-generated CAR-T cells and priming antigen-specific T cell responses

Having established a functional T-cell-priming niche *in vivo*, we next evaluated whether iCTA could recruit and amplify *in vivo*-generated CAR-T cells. We employed CD5-targeted mRNA-LNPs encoding an anti-DLL3 CAR, which can interact with DLL3, an atypical Notch ligand highly expressed in small-cell lung cancer (SCLC).^29^ T-cell targeting of the LNPs was first evaluated in Ai9-tdTomato reporter mice (**Figure S2K**). After administration of LNP containing mRNA encoding Cre, evident tdTomato expression was observed in splenic T cells (**Figure S2L**), demonstrating efficient *in vivo* T-cell transfection.

Then, the impact of iCTA niche on CAR-T cells was evaluated. As shown in **Figure 2J**, LNPs were administered intravenously on day 0, followed two days later by subcutaneous injection of DLL3 antigen-loaded iCTA (iCTA^DLL3^). CAR⁺ T cells were subsequently monitored in peripheral blood and lymphoid organs. On days 3 and 5, flow cytometric analysis revealed the recruitment of CAR⁺ T cells (**Figure 2K**). In the dLNs, CAR⁺ T-cell frequencies increased to approximately 1.80- (*P* = 0.0034) and 1.84-fold (*P* = 0.0106) those of untreated controls on days 3 and 5, respectively (**Figure 2L**). A similar proliferative effect was also observed in the peripheral blood, where iCTA^DLL3^ treatment increased the abundance of CAR⁺ T cells by approximately 2.67- (*P* = 0.0003) and 2.13-fold (*P* = 0.0056), respectively, on day 3 and day 5 (**Figure 2M**). A similar trend was also observed among splenocytes (**Figure S2M**). Moreover, CAR tended to be more expressed in CD8^+^ T cells after the combination of iCTA^DLL3^ (**Figure 2N**), suggesting that iCTA supports the recruitment and sustained expansion of *in vivo*- generated CAR-T cells.

We next evaluated the ability of the iCTA platform to induce antigen-specific T-cell activation *in vivo*. As shown in **Figure 2O**, CD45.2⁺ OT-I T cells were isolated *ex vivo* and intravenously adoptively transferred into B6 CD45.1 recipient mice on day 0. This model allowed us to directly track antigen-specific T cell responses *in vivo*.^30^ Two days later, OVA- loaded iCTAs (iCTA^OVA^) were administered subcutaneously, followed by flow cytometric analysis of the transferred CD45.2⁺ OT-I cells. In mice receiving the combination of iCTA^OVA^ and OT-I cells, we observed recruitment of the transferred OT-I cells to the injection site (**Figure 2P**), together with expansion of OT-I cells in the dLNs (**Figures 2Q** and **2R**). Further analysis of CD45.2⁺ OT-I cells in the dLNs showed that iCTA^OVA^ pulse significantly increased the expression of both Ki67 and CD25 (**Figures 2S** and **2T**), demonstrating the potential of iCTA to induce antigen-specific T-cell activation *in vivo*. Together, these findings show that the iCTA niche can simultaneously recruit and amplify *in vivo*-generated CAR-T cells while priming antigen-specific T-cell responses.

### iCTA remodels the tumor microenvironment

Having established that iCTA promotes T-cell recruitment and priming, we next asked whether these responses translated into remodeling of the tumor microenvironment (TME). After two immunizations, we assessed intratumoral T-cell infiltration and activation (**Figure 3A**). ELISA of tumor supernatants showed that iCTA treatment increased IFNγ, IFNβ, and TNFα levels, indicating a more inflammatory, immune-supportive TME (**Figure 3B**). We then performed 10× scRNA-seq on CD45⁺ leukocytes from B16-OVA tumors, which identified myeloid, T cell, and NK cell lineages (**Figures 3C** and **3D**). Compared with Mn alone, iCTA increased the overall lymphocyte fraction, including T and NK cells, and raised intratumoral CD4⁺ and CD8⁺ T cell abundance (**Figures 3E** and **3F**). Flow cytometry further showed higher frequencies of IFNγ⁺ CD8⁺ T cells, IFNγ⁺ CD4⁺ T cells, and IFNγ⁺ NK cells (**Figure 3G**).

**Figure 3.**
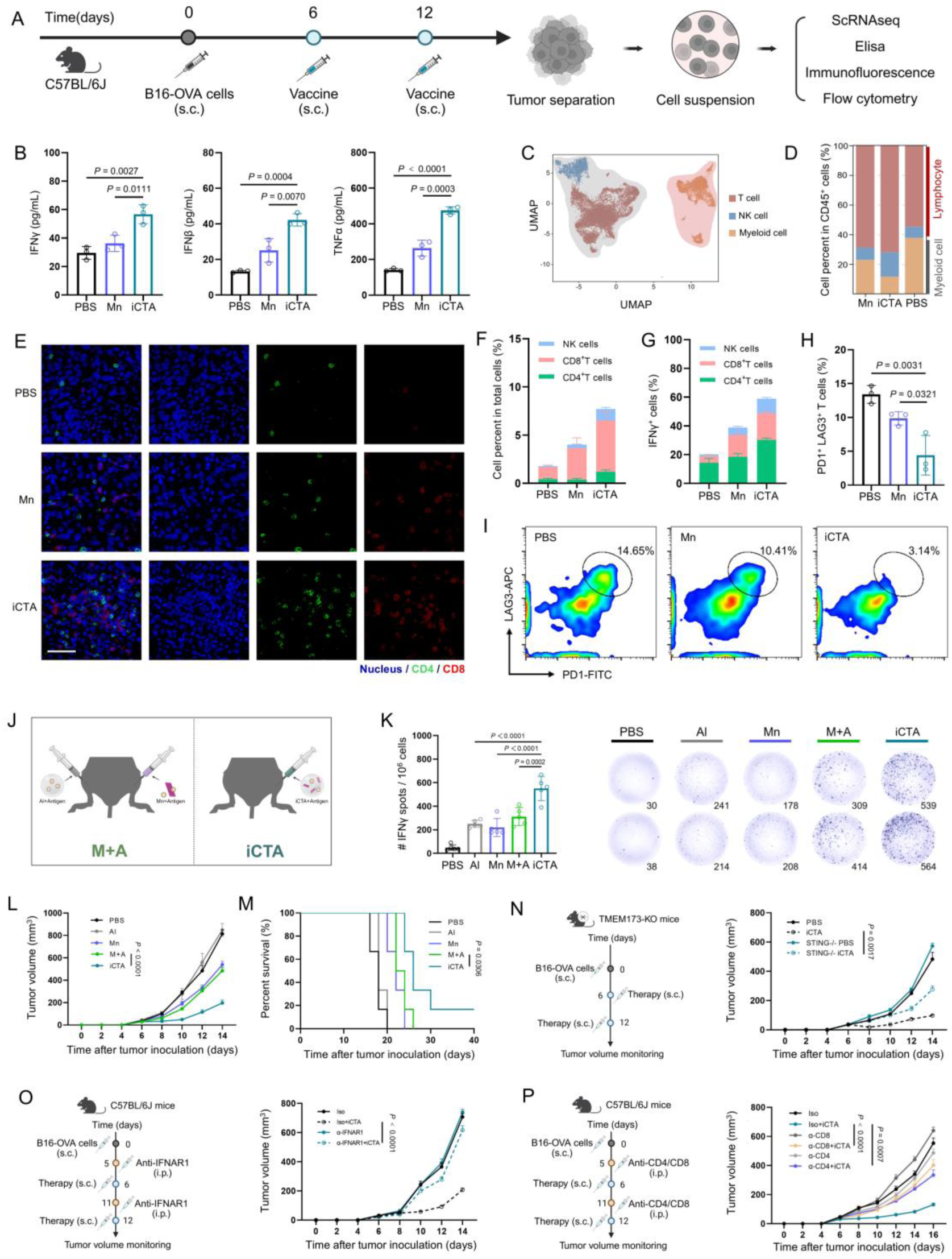
Evaluation of iCTA-mediated TME reshaping *in vivo*. (A) Schematic illustration of the experimental design for TME profiling after iCTA tumor vaccine treatment. B16-OVA tumors were established on day 0; mice were vaccinated on days 6 and 12; tumors were collected on day 14. One fraction was used for ELISA, immunofluorescence, and flow cytometry; CD45^+^ cells from the remaining fraction were enriched for 10× scRNA-seq. (B) ELISA quantification of IFNβ, IFNγ, and TNFα in tumor homogenates (n = 3). (C, D) scRNA-seq analysis of TME composition. (C) UMAP of major clusters and (D) relative cell- type abundance. (E) Representative immunofluorescence of intratumoral CD4^+^ T cells (green), CD8^+^ T cells (red), and nuclei (blue). Scale bar, 50 μm. (F) Frequencies of CD4^+^ T cells, CD8^+^ T cells, and NK cells in tumors by flow cytometry (n = 3). (G) IFNγ production by CD4^+^ T cells, CD8^+^ T cells, and NK cells in tumors (n = 3). (H, I) T-cell exhaustion in tumors. (H) quantification and (I) representative plots of PD1^+^LAG3^+^ T cells (n = 3). (J) Comparison of the antitumor efficacy between co-delivered iCTA and spatially separated administration (M+A). Left: schematic illustration of the injection method on days 6 and 12; right: tumor growth kinetics (n = 6). (K) IFNγ ELISPOT of tumor-infiltrating lymphocytes. Quantification and representative images (n= 5). (L) Tumor growth curves under indicated treatments (n = 6). (M) Survival from day 0 to day 40 after tumor inoculation (n = 6). (N) TMEM173-deficient model used to test whether the antitumor activity is STING-dependent. Left: treatment schedule; right: tumor growth kinetics (n = 5). (O) IFNAR1 blockade model used to evaluate whether the antitumor effect of iCTA is related to type I interferon. Left: anti-IFNAR1 on days 5 and 11, iCTA vaccine on days 6 and 12; right: tumor growth kinetics (n = 6). (P) CD4 or CD8 depletion study. Left: experimental design (anti-CD4 or anti-CD8 on days 5 and 11; iCTA vaccine on days 6 and 12). Right: tumor growth curves (n = 6). Data in (B), (F), (G), (H), and (K) are presented as the mean ± SD. Data in (L), (N), (O), and (P) are presented as the mean ± SEM. Statistical significance was calculated via one-way ANOVA with a Tukey post-hoc test (B, H, K, L, N, O, and P) or two-sided log-rank (Mantel-Cox) test (M).

As tumor-infiltrating T cells frequently acquire dysfunctional or exhausted states, we next examined their differentiation states. Sub-clustering identified nine T cell subsets,^31,32^ including a CD8Tex-*LAG3* population expressing high levels of *PDCD1*, *LAG3*, and *HAVCR2* (**Figures S3A-S3C**), whose frequency was obviously reduced (**Figure S3C**). Our flow cytometry analysis correspondingly showed the decrease of PD1^+^ LAG3^+^ T cells after iCTA-treated (**Figures 3H** and **3I**). These findings suggest that, in addition to increasing lymphocyte infiltration, iCTA may shift intratumoral T-cell states away from a highly exhausted phenotype, potentially creating a more permissive environment for *in vivo* CAR-T-cell function.

In addition to T cells, we further examined intratumoral myeloids and identified a distinct cDC1-*CCL22* subset (**Figures S3D-S3F**),^31^ whose abundance was markedly increased by iCTA (**Figure S3F**). This cluster exhibited elevated MHC-II signatures (**Figure S3G**),^33^ mirroring the MHC-II upregulation observed in injection site DCs. Consistently, iCTA increased the frequency of OVA₃₂₃₋₃₃₉ tetramer⁺ CD4⁺ T cells in tumor-draining lymph nodes (tdLN, **Figure S3H**), further supporting enhanced antigen-specific T-cell immunity.

Next, we comprehensively validated the biosafety of iCTA after twice injection *in vivo*. As shown in **Figure S3I**, the body weight of the mice in the iCTA group remained stable. Serum biochemistry (ALT, AST, BUN, and LDH) remained within normal ranges, and there was no significant difference compared with the healthy PBS group (**Figure S3J**). In addition to biochemical indicators, we also monitored the levels of representative cytokines (IFNγ, IL6, TNFα, and IL1β) in the serum, which were related to cytokine release syndrome (CRS) in the clinic. Compared with the PBS group, the iCTA group exhibited similar levels for all these representative cytokines (**Figure S3K**) and hematological parameters (**Table S1**). Limited necrosis, fibrosis, or inflammatory infiltration in the main organs (heart, liver, spleen, lungs, kidneys) were observed (**Figure S3L**).

### Spatially coupled Al–Mn adjuvanticity underlies the iCTA immune niche

Having established that iCTA forms a local T-cell-priming niche, recruits *in vivo*-generated CAR-T cells, mobilizes antigen-specific T-cell responses, and remodels the TME, we next defined the mechanistic requirements underlying these effects. We first asked whether the spatial integration of Al and Mn was necessary for iCTA activity. Compared with the co- formulated Al-Mn composite, administration of Al and Mn into contralateral hind limbs (M+A) reduced weakened tumor-infiltrating T-cell IFNγ responses (**Figures 3J** and **3K)** and tumor control relative to iCTA (**Figures 3L** and **3M**). These findings indicate that co-localization of Al-mediated antigen retention and Mn-mediated innate stimulation is a functional requirement of the iCTA niche, rather than the two components acting independently.

We next dissected the signaling pathway supporting this activity. iCTA inhibited tumor growth after two administrations, whereas this effect was markedly attenuated in STING- deficient *TMEM173*⁻/⁻ mice (**Figure 3N**) and following IFNAR1 blockade (**Figure 3O**), establishing the STING–type I IFN axis as a key mediator of iCTA adjuvanticity. Antibody- mediated depletion further showed that CD8⁺ T cells were required for tumor control, while CD4⁺ T-cell depletion also impaired therapeutic efficacy (**Figure 3P**), indicating that iCTA ultimately engages coordinated adaptive T-cell immunity.

Together, these findings indicated that Al retains and presents antigen locally, Mn provides STING-dependent innate activation, and their spatial coupling creates an immune niche that links APC activation with both CAR-T support and endogenous T-cell immunity. This mechanistic framework explains how the local niche characterized above can propagate from the injection site to draining lymphoid tissues and the tumor, thereby supporting *in vivo*- generated CAR-T cells while broadening antitumor immunity.

### iCTA potentiates adoptively transferred antigen-specific T cells

The increased priming of T cells and the improved programming of TME motivated us to further evaluate the performance of iCTA combined with adoptive cell transfer therapy (ACT). Given the antigen-specific activation of iCTA, we used ACT related to T cell receptors. In the B16-OVA tumor model, mice received two infusions of either OT-I or OT-II cells in combination with iCTA^OVA^ (**Figure 4A**). OT-I combined with iCTA^OVA^ produced pronounced tumor growth inhibition compared with OT-I therapy alone and significantly prolonged survival (**Figures 4B** and **4C**), indicating that the local priming niche can potentiate CD8⁺ cytotoxic T cell therapy. Notably, iCTA^OVA^ similarly increased the antitumor activity and extended survival (**Figures 4D** and **4E**), highlighting the contribution of antigen-specific CD4⁺ T-cell responses to sustained antitumor immunity.

**Figure 4.**
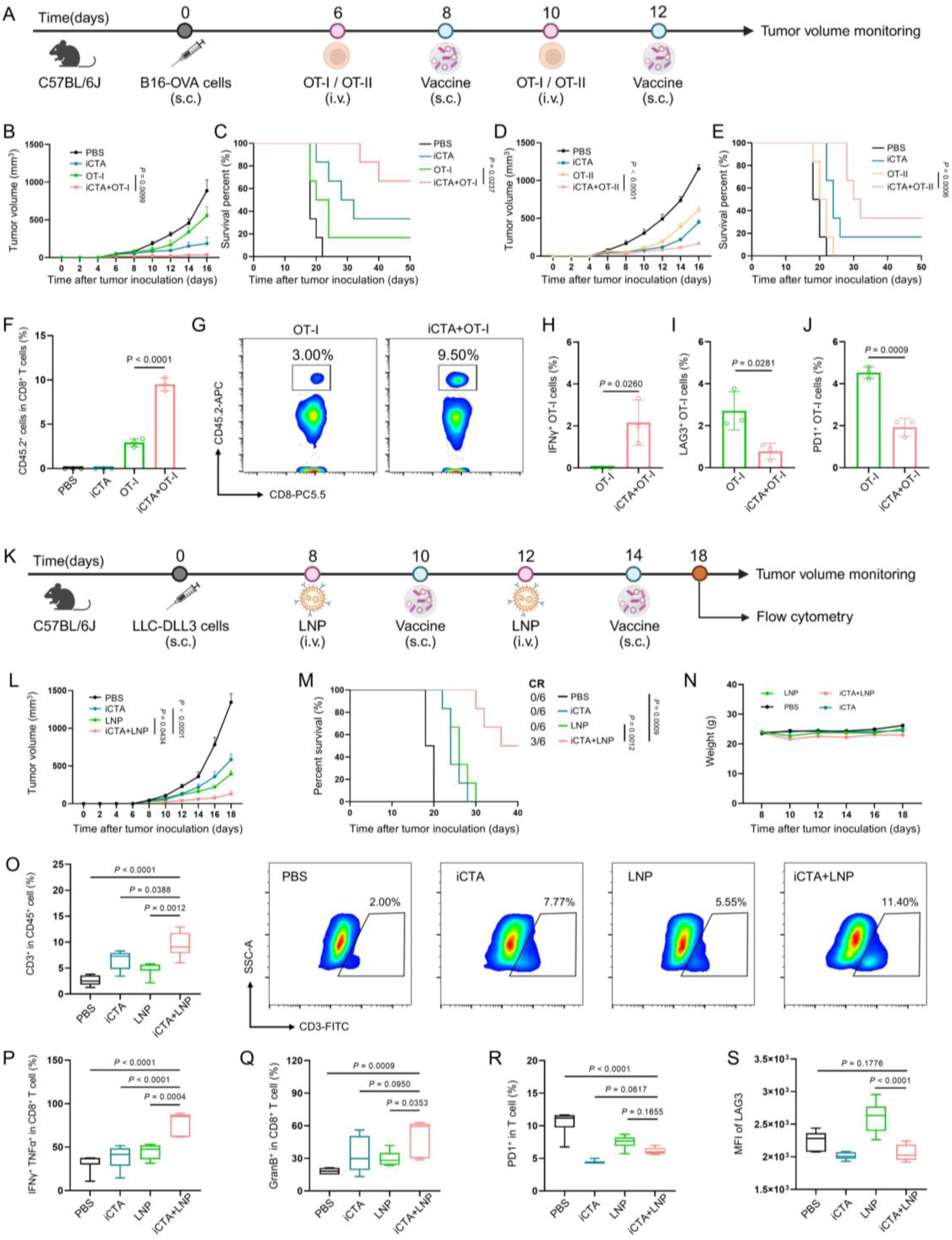
Evaluation of therapeutic efficacy of combination therapy. (A) Experimental design for combining iCTA tumor vaccine with OT-I or OT-II transfer. C57BL/6J mice bearing B16-OVA tumors (day 0) received OT-I/OT-II cells intravenously (days 6 and 10) and iCTA^OVA^ subcutaneously (days 8 and 12). Tumors were measured every 2 days; samples were collected on day 14 for OT-I analysis. (B-E) Antitumor efficacy from different treatment groups. (B, D) Tumor growth and (C, E) survival (n = 6). (F, G) Intratumoral CD45.2^+^ OT-I infiltration by flow cytometry. (F) Quantification and (G) representative plots (n = 3). (H) Activation of intratumoral CD45.2^+^ OT-I cells assessed by IFNγ expression (n = 3). (I, J) Exhaustion of CD45.2^+^ OT-I cells in tumors. Proportions of (I) LAG3^+^ and (J) PD1^+^ cells (n= 3). (K) Experimental design for combining iCTA tumor vaccine with LNP that used for *in vivo*- generated CAR-T cells in LLC-DLL3 tumors. LNPs were given intravenously (days 8 and 12); iCTA vaccine subcutaneously (days 10 and 14). Tumors were tracked every 2 days; tumors and spleens collected on day 18. (L) Tumor growth curves (day 0-18, n = 6). (M) Survival (day 0–40) and complete response (CR) counts (n = 6). (N) Body weight monitoring after treatment (days 8-18, n = 6). (O) Intratumoral T cell infiltration. Quantification and representative plots (n = 6). (P, Q) CD8^+^ T cell cytotoxicity in tumors. (P) IFNγ^+^ TNFα^+^ and (Q) GranB^+^ fractions (n = 6). (R, S) Splenic T cell exhaustion. (R) PD1^+^ frequency and (S) LAG3 MFI (n = 6). Data in (F), (H-J) and (O-S) are presented as the mean ± SD. Data in (B), (D), (L) and (N) are presented as the mean ± SEM. Statistical significance was calculated via one-way ANOVA with a Tukey post-hoc test (F and O-S), two-tailed Student’s *t* test (H-J), or two-sided log-rank (Mantel- Cox) test (C, E, and M).

We next tracked transferred CD45.2⁺ OT-I cells in CD45.1 recipient mice to determine how iCTA influenced their activation and tumor trafficking. In tdLNs, combination treatment increased the abundance of OT-I cells and elevated the proportion of IFNγ⁺ OT-I cells (**Figures S4A-S4D**), consistent with enhanced cytotoxic programming before tumor entry. In regard with the TME, iCTA^OVA^ increased intratumoral accumulation of adoptively transferred OT-I cells by approximately 3.0-fold (*P* < 0.0001), compared with OT-I adoptive transfer monotherapy (**Figures 4F** and **4G**). These infiltrated OT-I cells also exhibited significantly higher IFNγ expression (**Figure 4H**), demonstrating that iCTA^OVA^ not only improves OT-I cell trafficking into tumors but also enhances their effector function *in situ*. In parallel, exhaustion-associated markers, including LAG3 and PD1, were substantially reduced on intratumoral OT-I cells following iCTA^OVA^ treatment (**Figures 4I** and **4J**). These findings indicated that the combined strategy enhanced antitumor efficacy while improving intratumoral infiltration and function of adoptively transferred T cells, providing a mechanistic rationale for its combination with in vivo-generated CAR-T cells.

### iCTA potentiates *in vivo* CAR-T therapy against established tumors

Encouraged by the above results, we next evaluated the adjuvanticity of iCTA in an *in vivo* anti- DLL3 CAR-T strategy. In the subcutaneous LLC-DLL3 lung cancer model, anti-DLL3- encoding LNPs were administered intravenously on days 8 and 12 to induce CAR-T cells *in vivo*, followed two days later by subcutaneous immunization with iCTA^DLL3^ to provide local antigenic and restimulation (**Figure 4K**). Compared with either monotherapy, the combination of *in vivo* CAR-T therapy and iCTA^DLL3^ effectively suppressed tumor progression (**Figures 4L** and **S4E**). Although LNP or iCTA^DLL3^ treatment alone modestly prolonged survival, the combination produced the most pronounced survival benefit (**Figure 4M**). Importantly, combination therapy did not induce obvious body weight loss throughout treatment (**Figure 4N**), supporting its preliminary safety.

We next characterized the immune remodeling induced by this combinational regimen. Flow cytometric analysis of tumor-infiltrating immune cells showed that combination therapy significantly increased the proportion of T cells within the TME compared with LNP group (**Figure 4O**). In addition, NK cell infiltration was markedly higher than that observed with either monotherapy (**Figure S4F**). Functional profiling further revealed a significant expansion of IFNγ⁺ TNFα⁺ cytotoxic CD8⁺ T cells in the combination group (**Figure 4P**), while iCTA^DLL3^ co-treatment further enhanced granzyme B expression in CD8⁺ T cells (**Figure 4Q**). These data demonstrate that iCTA not only increases effector-cell abundance in tumors but also strengthens their cytolytic activity.

We further examined whether iCTA influenced systemic T cell dysfunction. In the spleen, the frequencies of PD1⁺ T cells were lower in the combination group than in PBS controls (**Figure 4R**). Combination therapy also reduced CTLA4⁺ T cells compared with CAR-T monotherapy (**Figure S4G**), and the mean fluorescence intensity of LAG3 on T cells was significantly diminished (**Figure 4S**). These findings indicate that iCTA helps restrain exhaustion-associated programs during *in vivo* CAR-T therapy, supporting more durable T cell functionality.

Collectively, iCTA addresses key constraints of transient *in vivo* CAR-T therapy by providing localized antigenic restimulation, increasing CAR-T-cell responses and tumor infiltration, limiting exhaustion-associated states, and mobilizing endogenous T- and NK-cell immunity. By coupling controllable CAR-T generation with a spatially confined immune-priming niche, iCTA increases the magnitude and functional persistence of the antitumor response, enabling improved control of established tumors.

## DISCUSSION

*In vivo* CAR-T engineering enables direct and transient programming of endogenous T cells, bypassing *ex vivo* manufacturing and permanent genomic modification and thereby offering a faster and potentially more controllable form of cell therapy. However, this transient nature may also limit the magnitude and functional persistence of CAR-T responses required to eradicate established tumors. Current strategies have largely sought to intensify T-cell activation *in situ*, for example through immunostimulatory lipid nanoparticles or co-delivery of cytokine- encoding mRNAs.^14,34,35^ Drawing from *in vitro* T cells strategies, excessive or premature stimulation may accelerate T-cell dysfunction within the TME.^10–12^ Previous attempts to boost the transferred CAR-T cells demonstrated the feasibility of vaccine-based boosting of adoptively transferred CAR-T cells.^26,36^ Amphiphile CAR-ligand vaccines, for example, exploit albumin-mediated trafficking to draining lymph nodes, where cognate antigen is displayed to stimulate pre-existing CAR-T cells and enhance antitumor activity.^26^ *In vivo* CAR-T therapy, however, creates a distinct opportunity: CAR-positive T cells are generated transiently within an otherwise intact immune system that remains available for therapeutic modulation. Thus, antitumor activity need not be confined to CAR-expressing cells; the much larger CAR- negative T-cell compartment, together with innate immune cells, can also be mobilized to support tumor control and remodel the TME. As a proof-of-concept, we developed an *in vivo* CAR-T adjuvant (iCTA) that builds on this principle but shifts the paradigm from direct lymphoid boosting of pre-existing CAR-T cells toward a local immune niche supporting *in vivo*-generated CAR-T cells together with the surrounding endogenous immune repertoire.

A single subcutaneous iCTA administration increased the abundance of CAR-positive T cells in draining lymph nodes and peripheral blood (**Figures 2L** and **2M**), extending the principle of vaccine-mediated CAR-T boosting to *in vivo*-generated CAR-T cells. iCTA also activated broader immune compartments, including engaged antigen-specific CD4⁺ and CD8⁺ T cells through antigen presentation via APCs (**Figures 2P-2T**). These effects further extended from the injection-site niche to the tumor, increasing T- and NK-cell infiltration, establishing an IFNγ-enriched microenvironment, and reducing exhaustion-associated T-cell states (**Figure 3A-3I**). Adoptive-transfer experiments further demonstrated enhanced trafficking and effector function of antigen-specific T cells within tumors (**Figures 4A-4J**). Thus, iCTA extends CAR- T boosting beyond the engineered CAR-positive population, harnessing the immune system to support *in vivo*-generated CAR-T cells while broadening endogenous antitumor immunity.

The adjuvanticity of iCTA depends on the spatial coupling of antigen exposure and innate immune stimulation. Alum provides local antigen retention, whereas Mn contributes STING- type I IFN signaling. Separating Al and Mn reduced T-cell responses and antitumor efficacy, while STING deficiency or IFNAR1 blockade impaired iCTA activity (**Figures 1N** and **1O**). This indicated that iCTA is not simply the sum of two adjuvants; rather, their co-localization integrates accessible antigen, innate activation, APC maturation, and costimulation within a shared immune niche. Consistent with this model, iCTA retained antigen locally, promoted DC- T-cell interactions, and enhanced both CAR-T-cell abundance and conventional antigen- specific T-cell responses (**Figures 1N-1P** and **2E-2G**).

This adjuvant effects translated into improved control of established tumors when iCTA was combined with transient anti-DLL3 CAR mRNA-LNP treatment (**Figures 4L-4N**). The combination increased tumor infiltration and cytotoxic activity while reducing exhaustion- associated phenotypes relative to CAR-T therapy alone (**Fig. 4O-4S**). These findings suggested that rather than prolonging CAR expression itself, a spatially restricted and independently tunable immune niche can provide antigenic restimulation, complementary endogenous immunity, and a more permissive TME. This strategy may therefore preserve the controllability associated with transient *in vivo* CAR-T generation while compensating for insufficient stimulation and durability.

Engaging the CAR-negative immune compartment could also reduce reliance on repeated CAR mRNA-LNP dosing. Broader adaptive immune recruitment may lower the dose or frequency of antibody-targeted nanoparticles, limiting T-cell perturbation from repeated or excessive CD3-, CD5-, or CD8-directed delivery. From a translational perspective, iCTA offers several features favorable for clinical implementation. Aluminum-based adjuvants have a decades-long clinical safety record, manganese supplementation operates within well- characterized pharmacological limits, and future iCTA synthesis uses a straightforward co- precipitation process amenable to large-scale GMP manufacturing. More broadly, the Al-Mn formulation described here represents one implementation of the iCTA concept rather than its only possible form. Other antigen-retaining and immunostimulatory platforms could similarly be engineered to couple transient *in vivo* CAR-T generation with endogenous adaptive immunity, providing a general framework for improving the efficacy of *in vivo* cellular immunotherapy.

### Limitations of the study

Several mechanistic questions remain unresolved, including the relative contributions of direct CAR engagement and MHC-dependent antigen presentation, whether APCs form a distinct functional synapse with CAR-T cells, and how CAR-positive and CAR-negative T-cell populations quantitatively contribute to tumor control. Beyond these mechanistic uncertainties, the preclinical scope of our study is still limited. Efficacy was evaluated exclusively in syngeneic murine tumor models bearing a single defined antigen, and the current work focused on one tumor type (LLC-DLL3). Future validation in humanized mouse models or patient- derived xenografts with heterogeneous antigen expression profiles, together with longitudinal immune monitoring and broader tumor testing, may further demonstrate the durability and translational scope.

## METHODS

### Mouse models

Female BALB/c, C57BL/6, TMEM173-KO, OT-I, OT-II, and B6-CD45.1 mice aged 6-8 weeks were purchased from the respective suppliers and housed under standard controlled conditions with ad libitum access to food and water. The animal facility was maintained at 23 °C with 55 ± 5% humidity on a 12 h light/12 h dark cycle. All animal procedures were approved by the Institutional Animal Care and Use Committees of the Institute of Process Engineering, Chinese Academy of Sciences (approval no. IPEAECA2023056).

### Preparation and characterization of iCTA

For preparation of iCTA, stock solutions of AlCl₃·6H₂O (185 mM) and MnCl₂·4H₂O (91 mM) were combined immediately before use. The reactor contained 80 mM Na₂HPO₄ at room temperature and stirred at 600 rpm. After electrode calibration, the initial pH was recorded and was expected to be approximately 9.0-9.5, and it was adjusted to the set-point of 9.2 ± 0.1. The metal feed was added by syringe pump at 0.50 mL/min. Standardized, carbonate-free 0.500 M NaOH was added by pH-stat feedback to maintain the set-point, with both inlets positioned in well-mixed regions to avoid contact between concentrated streams. After the feed, the suspension was stirred for 2 h and bath-sonicated for 10 min. Particles were collected by centrifugation (6000 × g, 15 min) and washed three times with PBS (pH 7.4). For size and charge analysis, iCTA was dispersed in PBS, and the size and zeta potential were detected using dynamic light scattering system (Malvern ZEN 3600 Zetasizer).

For transmission electron microscopy (TEM), freshly prepared iCTA suspension was dropped onto carbon-coated copper grids (Electron Microscopy China) and allowed to adsorb. Excess liquid was removed with filter paper, and the grids were negatively stained with uranyl acetate. After washing and air-drying, samples were imaged by TEM (FEI tecnai F20), operated at 120 kV. TEM Mapping and EDS analysis within the iCTA formulation were further performed by energy-dispersive X-ray spectroscopy (Oxford X-MAX 80T) coupled to TEM. For XPS analysis, iCTA samples were lyophilized to obtain dry powders. To assess whether the valence state of manganese has changed, we also conducted XPS testing on the manganese adjuvant. The dried samples were mounted onto conductive adhesive substrates and then operated on an ESCALAB 250Xi instrument (Thermo Fisher Scientific). For XRD analysis, iCTA was lyophilized and then operated on Empyrean high-resolution powder X-ray diffractometer (PANalytical B.V.).

#### Antitumor effect and *in vivo* antibody blockade

For therapeutic tumor vaccination, C57BL/6J mice were subcutaneously inoculated with 5 × 10^5^ B16-OVA tumor cells on day 0. Mice were randomized when tumors became palpable and immunized subcutaneously with OVA antigen (10 μg per mice) formulated with adjuvants on days 6 and 12. For the spatially separated M+A treatment, alum and manganese adjuvant- containing formulations were injected at distinct subcutaneous sites. Among them, the adjuvant dose was the same for the two injection routes: 20 μg per mouse for alum adjuvant and 20 μg per mouse for manganese adjuvant. Tumor size was measured every 2 days with a digital caliper, and tumor volume was calculated as length × width² / 2. For *in vivo* type I interferon receptor blockade, mice bearing B16-OVA tumors received intraperitoneal injections of 200 μg (in 100 µl PBS) anti-IFNAR1 antibody before vaccination. Anti-IFNAR1 antibody was administered on days 5 and 11, followed by subcutaneous iCTA tumor vaccination on days 6 and 12. Control mice received an equal amount of isotype control antibody. Tumor growth was monitored every 2 days. To assess the requirement for STING signaling, TMEM173-deficient mice and corresponding wild-type control mice were subcutaneously implanted with B16-OVA tumor cells on day 0. Mice were treated with the iCTA tumor vaccine on days 6 and 12, and tumor growth was recorded every 2 days. For T cell blockade experiments, tumor-bearing mice received intraperitoneal injections of 200 μg (in 100 µl PBS) anti-CD4 or anti-CD8 antibody on days 5 and 11. iCTA tumor vaccine was administered subcutaneously on days 6 and 12. Control animals received isotype control antibody. Tumor volumes were measured every 2 days to evaluate the contribution of CD4⁺ and CD8⁺ T cells to iCTA vaccine-mediated antitumor efficacy.

#### Assessment of antigen display

For the evaluation of plasma membrane anchoring, DC2.4 cells were seeded onto sterile glass coverslips and allowed to adhere overnight. Cells were then incubated with iCTA for 6 h. After incubation, samples were gently washed with PBS three times and fixed with 2.5% glutaraldehyde for 10 min. The cells were subsequently dehydrated through a graded ethanol series, dried by critical-point drying, and sputter-coated with gold. Cell morphology and iCTA- cell interactions were examined using scanning electron microscopy (JSM-6700F).

For confocal laser scanning microscopy observation of antigen display, iCTA was first labeled with lumogallion and then incubated with Cy5-labeled OVA for 2 h to allow antigen adsorption. After washing three times, the OVA-adsorbed iCTA were added for incubating with DC2.4 cells for 6 h. For CD19 protein imaging, lumogallion-labeled iCTA was mixed with Alexa Fluor 647-labeled CD19 protein, and the antigen-adsorbed iCTA was incubated with hBMDCs for 6 h. After incubation, cells were washed with PBS three times, fixed with paraformaldehyde, and stained with DAPI to visualize nuclei. Confocal laser scanning microscope (Leica) was performed to assess the antigen display on the cell surface.

#### Antigen accessibility analysis

To evaluate the surface accessibility of iCTA, Alexa Fluor 647-labeled CD19 antigen was mixed with iCTA, and the antigen-adsorbed iCTA was incubated with hBMDCs for 6 h. Then cells were further stained with the anti-CD19 antibody as described above. After blocking with 3% BSA for 2 h, cells were washed three times with PBS. Subsequently, anti-CD19 antibody diluted 100-fold in 3% BSA/PBS solution was added, and cells were incubated overnight at 4°C, followed by three washes with PBS. AF488-conjugated secondary antibody diluted 1:200 was then added, and cells were stained for 2 h at room temperature. After three washes with PBS, cells were stained with DAPI for 10 min. hBMDCs were immediately imaged using a confocal laser scanning microscope (Leica) with LAS AF Lite 2.6.0 software.

QCM-D (Biolin Scientific/Q-Sense, Sweden) was employed to evaluate the interaction between CD19-adsorbed iCTA and the anti-CD19 antibody. Chips were modified by spin- coating iCTA or CD19-adsorbed iCTA, respectively. After mounting the modified chips, iCTA (0.5 mg/mL) was introduced into the QCM chamber at a continuous flow rate of 50 μL/min. The change in vibrational frequency (ΔF) was measured to characterize the adsorption behavior of anti-CD19.

### Evaluation of APC activation *in vitro*

Bone marrow cells were isolated from mouse femurs and tibias under sterile conditions. After red blood cell lysis, cells were cultured in complete medium supplemented with 10 ng/mL granulocyte-macrophage colony-stimulating factor (GM-CSF) and 20 ng/mL interleukin 4 (IL4) or 20 ng/mL macrophage colony-stimulating factor (M-CSF) to BMDCs or BMDMs. Non- adherent or loosely adherent BMDCs and adherent BMDMs were collected after differentiation and used for subsequent stimulation assays.

For western blot analysis, cells were stimulated with the indicated adjuvants for 24h. Next, cells were lysed in protein extraction buffer containing protease and phosphatase inhibitors. Equal amounts of protein were separated by SDS-PAGE and transferred to PVDF membranes. Membranes were blocked and incubated with primary antibodies against GAPDH, phospho- IRF3, and phospho-STING, followed by HRP-conjugated secondary antibodies. GADPH was used as a housekeeping protein to normalize the levels of protein. Protein bands were visualized using chemiluminescence detection.

For ELISA analysis of cytokine secretion, cell culture supernatants were collected and clarified by centrifugation after stimulation for 24h. Cytokine secretion levels were measured using ELISA kits according to the manufacturers’ protocols. Absorbance was recorded using a microplate reader, and cytokine concentrations were calculated from standard curves.

For the co-stimulation and antigen-presentation evaluation, BMDCs were stained with anti-mouse CD11c, CD40, CD86, MHC-I, MHC-II, and H2-K^B^-OVA_257-264_ antibodies at 4 ℃ for 30 min after stimulation with indicated adjuvants. After three washes, the cells were analysed by a flow cytometry system (Beckman) using CytExpert Pro 2.3.

#### Establishment of human CD19-CAR T cells *in vitro*

LNPs were formulated at an N:P ratio of 4.5 based on established ionizable lipid formulations. Using microfluidic mixing, an aqueous mRNA solution (in 10 mM citrate buffer, pH 4) was combined with an ethanolic lipid phase, which containing ALC0315, DSPC, cholesterol, and DMG-PEG2000, at a 3:1 aqueous-to-ethanol ratio. For antibody-conjugated LNPs, 0.5% of the PEG-lipid was replaced with DSPE-PEG2000-maleimide to enable subsequent bioconjugation. Antibody conjugation was performed via SATA–maleimide coupling chemistry. Purified rat anti-mouse CD5 antibody was thiolated using SATA according to the manufacturer’s protocol, followed by hydroxylamine-mediated deacetylation to expose free sulfhydryl groups. The thiolated antibody was then incubated with maleimide-functionalized LNPs at a defined molar ratio for 1-2 h under gentle agitation, allowing covalent thioether bond formation. Unconjugated antibody was removed by ultrafiltration, and the resulting antibody-LNP conjugates were buffer-exchanged into PBS and stored at 4°C until use.

Human PBMCs were isolated from healthy donors by density gradient centrifugation according to the manufacturer’s instructions. Isolated PBMCs were pre-incubated with Dynabeads Human T-Activator CD3/CD28 for 24 h, after which the Dynabeads were removed, and T cells were washed twice with serum-free RPMI 1640 medium for subsequent transfection. PBMCs were resuspended in serum-free medium and seeded into 96-well plates at a density of 1.0×10⁵ cells per well. CD5-conjugated LNPs loaded with CD19 CAR-encoding mRNA (0.5 μg mRNA per well) were then added, and cells were transfected for 24 h.

#### Evaluation of T cell activation *in vitro*

For CAR-T cell activation, hBMDCs were incubated with CD19 antigen formulated with the indicated adjuvants for 24 h. CD19 CAR-T cells were then added and co-cultured with antigen- loaded DCs at a cell ratio of 1:1 for 48 h. CAR-T cell activation was assessed by flow cytometry using anti-human CD3 and Ki67 antibodies.

Single-cell suspensions were prepared from spleens of OT-I transgenic mice by mechanical dissociation followed by red blood cell lysis. CD8⁺ T cells were purified by negative selection using a magnetic bead-based isolation kit (Biolegend) according to the manufacturer’s instructions. Purified OT-I cells were used immediately for co-culture assays. For OT-I activation, BMDCs were stimulated with OVA antigen formulated with the indicated adjuvants for 24 h. After stimulation, purified OT-I CD8⁺ T cells were added to the BMDC cultures and co-cultured for an additional 48 h. T cell activation and proliferation were analyzed by flow cytometry using mouse antibodies against CD3, CD8, CD25, Ki67 and IFNγ.

#### Characterization of the injection site niche

Indicated adjuvants were incubated with APC-labeled OVA to allow antigen adsorption. The antigen-adsorbed adjuvants were administered subcutaneously. *In vivo* fluorescence imaging was performed at 0, 12, 24, 48, and 168 h after injection using the animal imaging system. Fluorescence signals at the injection site were quantified using identical imaging parameters across groups. Subcutaneous injection site tissues were excised at 24h after adjuvant administration. Samples were fixed in glutaraldehyde, washed, dehydrated through graded ethanol, and dried. After sputter coating with a conductive metal layer, the microstructure of the injection-site nodule was examined by SEM. For H&E staining, tissue sections were deparaffinized, rehydrated, and stained with hematoxylin and eosin following standard procedures. Histopathological changes at the injection site were imaged using bright-field microscopy. For immunofluorescence staining, injection site tissues were fixed, paraffin- embedded, and sectioned. After deparaffinization, rehydration, and antigen retrieval, sections were blocked and incubated with primary antibodies against CD3 and CD11c. Fluorescent secondary antibodies were then applied, and nuclei were counterstained with DAPI. Images were acquired by confocal laser scanning microscope (Leica) to evaluate the spatial distribution of T cells and DCs at the injection site.

#### Analysis of injection site immune cells

Subcutaneous tissues together with adjacent muscle tissues surrounding the injection site were harvested and minced into small fragments. Tissues were enzymatically digested with collagenase-II-containing digestion buffer to generate single-cell suspensions. After filtration through a cell strainer and red blood cell lysis, cells were stained with a viability dye and fluorophore-conjugated mouse antibodies against CD45, CD3, CD11c, H2-K^B^-OVA_257-264_, MHC-II, CD40, CD80, and CD86. Samples were analysed by a flow cytometry system (Beckman) using CytExpert Pro 2.3.

Single-cell suspensions from injection-site tissues were prepared as described above, and CD11c⁺ dendritic cells were isolated by magnetic-activated cell sorting. Purified cells were collected for bulk RNA sequencing. Total RNA was extracted using TRIzol reagent, and RNA integrity was evaluated with an Agilent 2100 Bioanalyzer and RNase-free agarose gel electrophoresis. Polyadenylated mRNA was enriched with Oligo(dT) beads, fragmented, and reverse-transcribed into cDNA using the NEBNext Ultra RNA Library Prep Kit for Illumina. Double-stranded cDNA was end-repaired, A-tailed, ligated with Illumina adapters, purified with AMPure XP Beads, and PCR-amplified to generate sequencing libraries. Libraries were sequenced on an Illumina NovaSeq 6000 platform by Gene Denovo Biotechnology Co., Ltd. (Guangzhou, China). Raw reads were processed with fastp (v0.18.0) for quality control, followed by removal of rRNA-derived reads. High-quality paired-end reads were aligned to the reference genome using HISAT2 (v2.2.4) with default settings. Transcript assembly and expression profiling were subsequently performed using StringTie (v1.3.1) based on the reference genome annotation.

Injection-site tissues were harvested at 24h after treatment and dissociated into single-cell suspensions as described above. Samples were prepared from 6 mice per group. CD45⁺ immune cells were enriched using CD45^+^ magnetic bead kit according to the manufacturer’s protocol. Following enrichment, cells were washed, counted, and assessed for viability using Dead Cell Removal Kit. Samples with a viability of at least 95% were used for single-cell library preparation. The Single-Cell 3′ Library Kit v3 from 10x Genomics was used for single-cell transcriptome amplification and library construction according to the manufacturer’s instructions and previously established protocols. Sorted single-cell suspensions were loaded onto a 10x Genomics microfluidic chip to generate cDNA libraries, which were then prepared and sequenced on an Illumina NovaSeq 6000 platform across six lanes. Gene expression libraries were generated using the CellRanger count pipeline with the STAR algorithm (10x Genomics, version 4), as described previously. The resulting count matrices were analyzed with Seurat R package version 3.1.4, retaining only genes detected in at least 10 cells per sample. Cells of poor quality were excluded based on UMI counts, the number of detected genes, and mitochondrial transcript content. Potential doublets were identified and removed using DoubletFinder in R. The remaining high-quality single-cell datasets were integrated with the Harmony package and normalized by total UMI counts, followed by scaling with regression against total UMI counts and mitochondrial gene percentage using a scale factor of 1 × 10^4^. UMAP was applied for dimensionality reduction and visualization. Downstream bioinformatic analyses were performed using Omicsmart, an interactive online platform for real-time data analysis (http://www.omicsmart.com).

#### Monitoring of *in vivo*-generated CAR-T cells

CD5-conjugated LNPs encapsulating DLL3 encoded mRNA were administered i.v. to wild- type C57 at doses of 1 mg/kg of mRNA on day 0. Next, mice received subcutaneous vaccination with the iCTA tumor vaccine containing DLL3 antigen on day 2. Mice were sacrificed after 0 day, 1 day, 3 days, and 5 days administration of LNP. Injection-site tissues, peripheral blood, spleens, and lymph nodes were collected at the indicated time points and processed separately to obtain single-cell suspensions for flow cytometric analysis. Single-cell suspensions from injection-site tissues were prepared as described above. For suspension preparation of peripheral blood, peripheral blood was collected into anticoagulant-containing tubes. Red blood cells were removed using red blood cell lysis buffer at room temperature for 10min. After lysis, the remaining leukocytes were washed with PBS and collected by centrifugation for subsequent flow cytometric staining.

Spleens were harvested and mechanically dissociated by gently grinding the tissues through a 70 μm cell strainer using the plunger of a syringe. The strainer was rinsed with PBS to maximize cell recovery, and the resulting cell suspension was collected by centrifugation. Red blood cells were subsequently removed using red blood cell lysis buffer at room temperature for 10min. After lysis, the remaining splenocytes were washed with PBS and resuspended in flow cytometry staining buffer for subsequent analysis. Lymph nodes were collected and mechanically dissociated by gently grinding the tissues through a 70 μm cell strainer using the plunger of a syringe. The strainer was rinsed with PBS, and the resulting single-cell suspension was collected by centrifugation. Cells were then washed and resuspended in flow cytometry staining buffer for subsequent antibody staining and flow cytometric analysis. Cells were stained with a viability dye and antibodies against CD45, CD3, CD4, CD8, CD25, and scFv.

### Activation of OT-I *in vivo*

CD45.2⁺ OT-I cells were intravenously transferred into CD45.1 B6 mice on day 0. Mice then received subcutaneous vaccination with the iCTA tumor vaccine on day 2. Injection-site tissues and draining lymph nodes were collected on day 4 for downstream analyses.

For immunofluorescence analysis, lymph nodes were harvested, embedded and sectioned. Sections were stained with an antibody against CD45.2 to visualize transferred OT-I cells, and nuclei were counterstained with DAPI. Images were acquired by confocal microscopy.

For flow cytometry, tissues were processed into single-cell suspensions according to above methods. Cells were stained with a viability dye and antibodies against CD45.2, CD45.1, CD8, CD25. For intracellular analysis, cells were fixed and permeabilized before staining for Ki67. Flow cytometry was used to evaluate the expansion and activation status of transferred OT-I cells.

#### Characterization of the tumor microenvironment

Tumor tissues were excised and enzymatically dissociated with collagenase I, collagenase IV and DNase to generate single-cell suspensions. After washing, tumor-infiltrating lymphocytes were enriched using a tumor-infiltrating lymphocyte isolation kit (Solarbio) according to the manufacturer’s protocol. Antigen-specific cytokine-producing T cells were quantified using IFNγ ELISPOT assays following the manufacturer’s instructions. Spots were developed, imaged and quantified with an ELISPOT reader.

For flow cytometry analysis of tumor-infiltrating immune cells, tumors were mechanically minced and enzymatically digested to obtain single-cell suspensions. After filtration and red blood cell lysis, cells were stained with a viability dye followed by fluorophore-conjugated antibodies against CD45, CD3, CD4, CD8, NK1.1, LAG3 and PD1. For intracellular cytokine detection, cells were fixed, permeabilized and stained for IFNγ. Samples were acquired by flow cytometry and analyzed using standard gating strategies.

For cytokine analysis in the TME, tumor tissues were collected, homogenized in cold lysis buffer or PBS containing protease inhibitors. Homogenates were centrifuged to remove debris, and the supernatants were collected. Concentrations of IFNβ, TNFα, and IFNγ were measured by ELISA according to the manufacturers’ protocols.

For single-cell RNA sequencing, tumor tissues were dissociated into single-cell suspensions as described above. CD45⁺ immune cells were enriched by magnetic bead-based sorting and processed for single-cell RNA sequencing using the 10× Genomics platform. The specific operation steps and data analysis methods are the same as injection-site.

#### Combination of adoptive cell transfer therapy and iCTA

Wild C57BL/6J mice were subcutaneously inoculated with B16-OVA tumor cells on day 0. OT- I or OT-II cells were intravenously transferred into tumor-bearing mice on day 6 and 10. Mice then received subcutaneous vaccination with the iCTA tumor vaccine on day 8 and 12. Tumor volume was measured every 2 days. Tumors and draining lymph nodes were collected on day 16 for downstream analyses. For flow cytometry, tissues were processed into single-cell suspensions. Cells were stained with a viability dye and antibodies against CD45.2, CD45.1, CD8, LAG3 and PD1. For intracellular analysis, cells were fixed and permeabilized before staining for IFNγ. Flow cytometry was used to evaluate the effector function and exhaustion status of transferred OT-I cells.

### Combination of *in vivo* CAR-T and iCTA

C57BL/6J mice were subcutaneously inoculated with 5 × 10^5^ LLC-DLL3 tumor cells suspended in 50 μL of sterile PBS or the indicated injection medium on day 0. Tumor cells were injected into the right flank region. Mice were monitored regularly for tumor establishment, and tumor-bearing mice were randomly assigned to the indicated treatment groups when tumors became palpable. Animals with unsuccessful tumor implantation or tumors outside the predefined size range before treatment were excluded according to prespecified criteria. For combination therapy, LNPs designed for in vivo generation of CAR-T cells were administered intravenously via the tail vein on days 8 and 12 in a total injection volume of 200 μL per mouse. The iCTA tumor vaccine was administered subcutaneously on days 10 and 14. For combination- treatment groups, mice received both LNPs and iCTA according to the schedules described above. Tumor growth was monitored every 2 days throughout the treatment period. The longest tumor diameter (length) and the perpendicular diameter (width) were measured using digital calipers, and tumor volume was calculated using the formula: tumor volume = length × width²/ 2. Body weight and general health status were monitored concurrently to assess treatment tolerability. Animals were euthanized at the predetermined experimental endpoint or when tumors reached the maximum size permitted by the approved animal protocol or when humane endpoint criteria were met. Tumors were collected at the indicated endpoint for subsequent analyses where applicable.

For immune profiling, tumors and spleens were harvested at the indicated time points and processed into single-cell suspensions. Cells were stained with a viability dye and antibodies against CD45, CD3, CD8, NK1.1, LAG3, PD1 and CTLA4. For intracellular effector molecule staining, cells were fixed, permeabilized and stained for IFNγ, TNFα and granzyme B. Flow cytometry was performed to assess T cell infiltration, cytotoxic function and exhaustion after combined iCTA vaccination and *in vivo* CAR-T treatment.

#### Safety evaluation

Major organs, including heart, liver, spleen, lung and kidney, were collected after treatment. For hematoxylin and eosin (H&E) staining, tissue sections were deparaffinized in xylene, rehydrated through decreasing concentrations of ethanol, and stained with hematoxylin and eosin using standard procedures. After dehydration and clearing, the sections were mounted and examined by light microscopy. Histopathological alterations associated with treatment were evaluated based on tissue architecture and the presence of pathological changes, including inflammatory-cell infiltration, cellular degeneration or necrosis, vascular abnormalities, edema, and other organ-specific lesions.

Peripheral blood was collected from mice and transferred into serum collection tubes without anticoagulant. Blood samples were allowed to clot at room temperature for approximately 30 min and subsequently centrifuged at 3,000 × g for 15 min. The resulting serum supernatant was carefully collected without disturbing the cellular fraction. Serum biochemical parameters were measured using an automated biochemical analyzer according to the manufacturer’s instructions. Alanine aminotransferase (ALT) and aspartate aminotransferase (AST) levels were determined as indicators of hepatocellular injury, whereas blood urea nitrogen (BUN) was measured to assess renal function. Lactate dehydrogenase (LDH) levels were additionally determined as a general indicator of systemic tissue injury.

For routine hematological analysis, peripheral whole blood was collected into tubes containing anticoagulant as an anticoagulant. Samples were gently mixed immediately after collection to prevent clot formation and were analyzed using an automated hematology analyzer according to the manufacturer’s instructions.

## QUANTIFICATION AND STATISTICAL ANALYSIS

Statistical analysis was performed by using GraphPad Prism 9.0.0. Statistical analysis was performed using two tailed Student’s t test (comparison between two groups) or one-way analysis of variance (ANOVA; comparison among three and more groups). Two-sided log-rank (Mantel-Cox) test was utilized to analyze the statistical significance of survival.

## AUTHOR CONTRIBUTIONS

Y.X. and Jy. C. conceived the idea. Y.X. and Y.S. designed the experiments. Y.S., Ys.M., T.G. and Y.Y. carried out the experiments. Y.S., T.G. and Yl.M. analyzed the data. J.C. and M.Q. designed the circRNA and the orthotopic DLL3-positive small-cell lung cancer model. M.Q. offered the engineered cancer cells. Y.L. and F.Y. offered *TMEM173* -/- mice model. Jy. C., G.M., X.H., F.Y. and M.Q. offered suggestion. Y.X. and Y.S. wrote the manuscript. Ys.M., Jy. C., T.G., Y.Y., and Yl.M. revised the manuscript. Y.X. provided funding and resources. All authors discussed the results and commented on the manuscript.

