## Supplementary material for "An adjuvant for in vivo CAR-T therapy potentiation against solid tumors": Figures S1-4 and Table S1

**SUPPLEMENTAL INFORMATION**

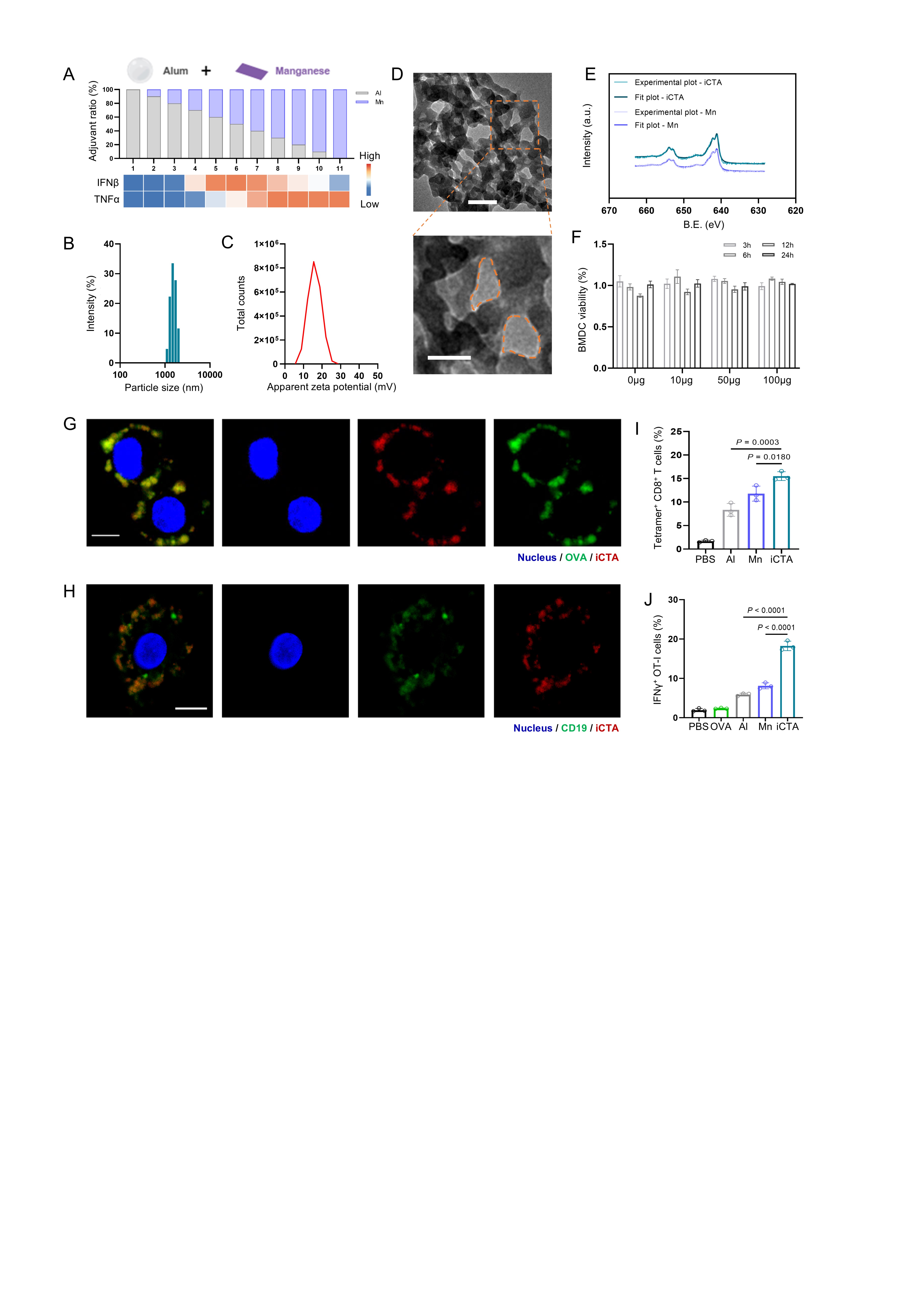

**Fig S1. Formulation and characterization of iCTA for T cell priming**

(A) Optimization of Al:Mn ratios. BMDMs were stimulated with the indicated formulations, and IFNβ and TNFα in supernatants were measured by ELISA at 24 h (n = 3).

(B, C) Particle size distribution (B) and surface charge (C) of iCTA measured by DLS.

(D) Representative TEM images of iCTA. Top scale bar, 200 nm; bottom scale bar, 100 nm.

(E) XPS spectra of Mn in manganese adjuvant and iCTA.

(F) Cell viability of BMDCs after co-incubation with iCTA, assessed by CCK8 assay (n = 3).

(G) Confocal microscopy images of DC2.4 cells after 6 h of incubation with labeled iCTA (red) loaded with Cy5-labeled OVA antigen (green). Scale bar, 5 μm.

(H) Confocal microscopy images of human bone marrow-derived dendritic cells (hBMDC) after 6 h of incubation with labeled iCTA particles (red) loaded with Alexa Fluor 647-labeled CD19 antigen (green). Scale bar, 5 μm.

(I) Antigen-specific T-cell priming by adjuvant-treated BMDCs co-cultured with mouse T cells (n=3).

(J) Flow cytometric analysis of intracellular IFNγ expression in OT-I cells (n = 3).

Data in (F), (I), and (J) are presented as the mean ± SD. Statistical significance was calculated via one-way ANOVA with a Tukey post-hoc test.

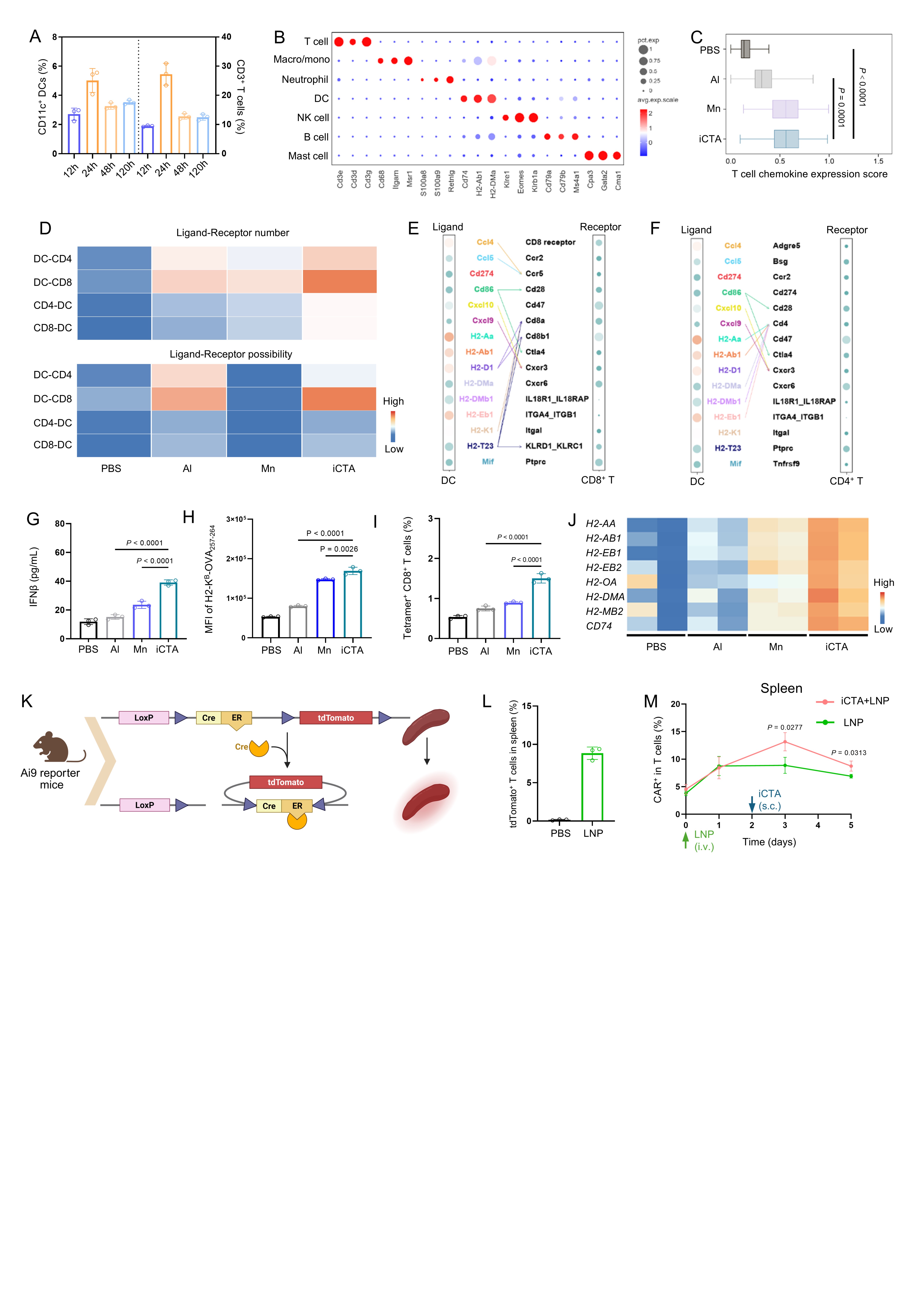

**Fig S2. Evaluation of APC-T interaction and T cell priming *in vivo***

(A) Flow cytometric quantification of the proportion of CD11c^+^ DCs and CD3⁺ T cells in local injection site tissue at 12, 24, 48, and 120 h after iCTA injection (n = 3).

(B) Bubble plot showing marker gene expression across clustered immune cell populations at the injection niche, identified by scRNAseq analysis 24 h after adjuvant administration.

(C) Gene set scoring analysis of the myeloid populations at the injection site. T cell chemokine expression scores (*CCL2, CCL3, CCL4, CCL5, CXCL9, CXCL10, CXCL11*) for the indicated adjuvants.

(D) Single-cell ligand–receptor interaction analysis between DCs and T cells. Heatmaps showing the number of ligand–receptor pairs and predicted interaction probabilities between DCs and T cells.

(E, F) Bubble plots of ligand–receptor combinations among the top 15 highly expressed pairs in DCs and T cells, depicting DC interactions with CD8^+^ T cells (E) and CD4^+^ T cells (F).

(G) Concentration of IFNβ in local injection site tissue at 24h after iCTA injection detected by ELISA (n = 3).

(H) Mean fluorescence intensity (MFI) of H2-K^B^-OVA on DCs surface in local injection site tissue at 24h after adjuvant injection detected by flow cytometry (n = 3).

(I) Flow cytometry analysis of the frequencies of H-2K^B^-SIINFEKL-tetramer^+^ CD8^+^ T cells after injection of the indicated adjuvants in the dLNs (n = 3).

(J) One day after injection of the indicated adjuvants, CD11c⁺ DCs were isolated from the injection site by magnetic-activated cell sorting and subjected to RNAseq. Heatmap depicting the expression of MHC-II related genes.

(K, L) Validation of the spleen-targeting capacity of CD5-conjugated LNPs using Ai9 reporter mice. (K) Schematic illustration of the Ai9 reporter mouse system. (L) Representative flow cytometry plots showing tdTomato⁺ T cells and quantification of their percentage among CD3⁺ T cells in the spleen (n = 3).

(M) Longitudinal frequencies of CAR^+^ T cells at spleen, comparing iCTA + LNP versus LNP alone (n = 3).

Data in (A), (C), (G-I), (L), and (M) are presented as the mean ± SD. Statistical significance was calculated via one-way ANOVA with a Tukey post-hoc test (C, G-I) or two-tailed Student’s *t* test (L, M).

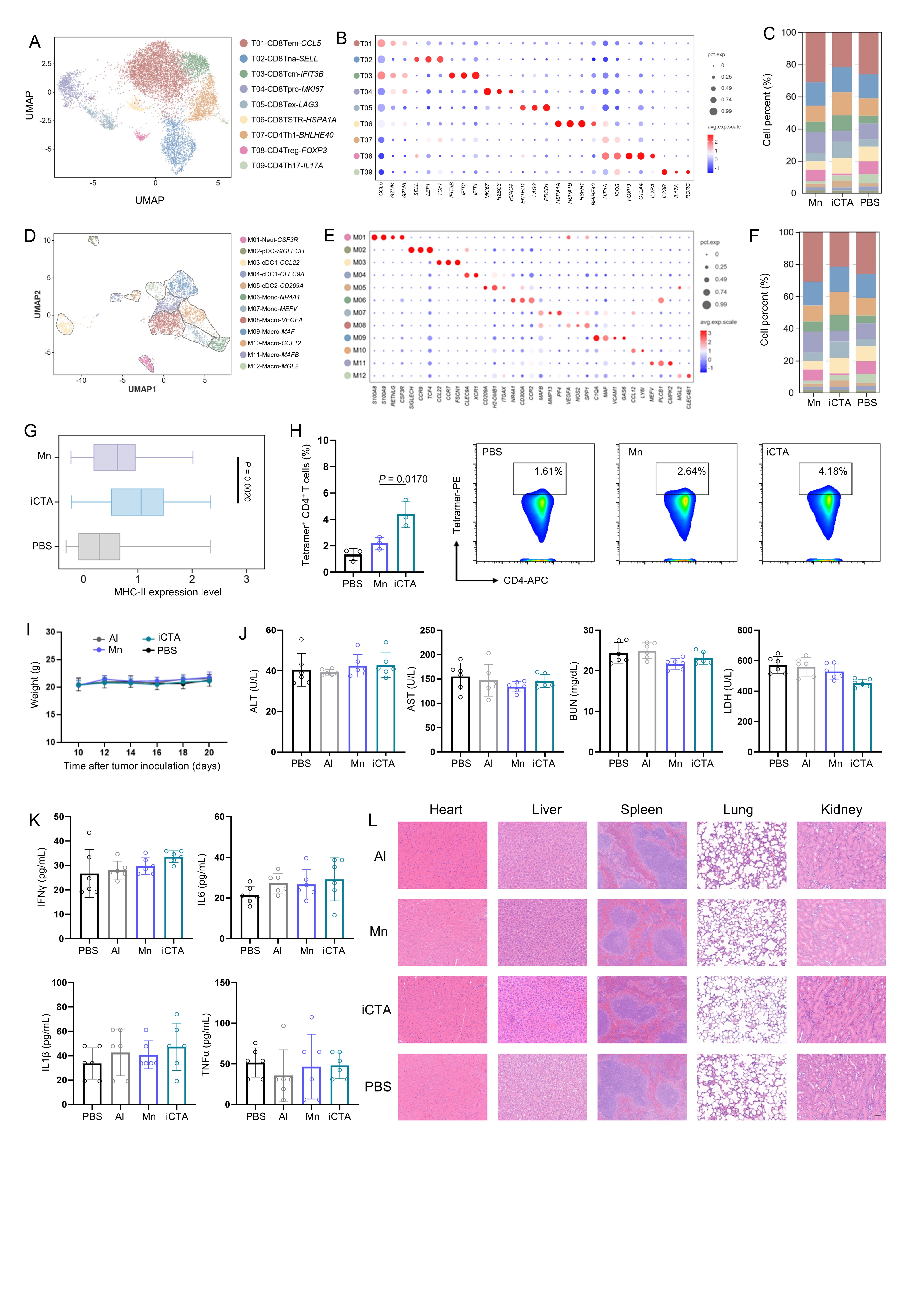

**Fig S3. Evaluation of the TME and biosafety**

(A-C) ScRNAseq re-clustering analysis of T cells in the TME. (A) UMAP map of different T cell subsets, (B) marker gene expression bubble plot, and (C) cell percent.

(D-F) Sub-clustering of myeloid cells, illustrated by UMAP plot (D), marker gene expression (E), and the proportional composition of myeloid cell subsets (F).

(G) Gene-set scoring of cDC1-*CCL22* population in the TME based on scRNAseq. MHC-II expression score across the indicated adjuvant groups.

(H) CD4^+^ antigen-specific T-cell priming by adjuvant-treated in tdLNs. Frequencies and representative plots of I-A^b^–ISQAVHAAHAEINEAGR tetramer^+^ CD4^+^ T cells (n = 3).

(I) Body weight curves of mice following administration of the indicated adjuvants (n = 6).

(J) Serum biochemical analyses of ALT, AST, LDH, and BUN at day 28 post-administration (n = 6).

(K) ELISA quantification of serum IFNγ, IL6, IL1β, and TNFα levels at day 28 post-injection (n = 6).

(L) Representative H&E-stained sections of heart, liver, spleen, lung, and kidney for histopathological evaluation. Scale bar, 50 μm.

Data in (G), (H), (J), and (K) are presented as the mean ± SD. Data in (I) is presented as the mean ± SEM. Statistical significance was calculated via one-way ANOVA with a Tukey post-hoc test (G, H).

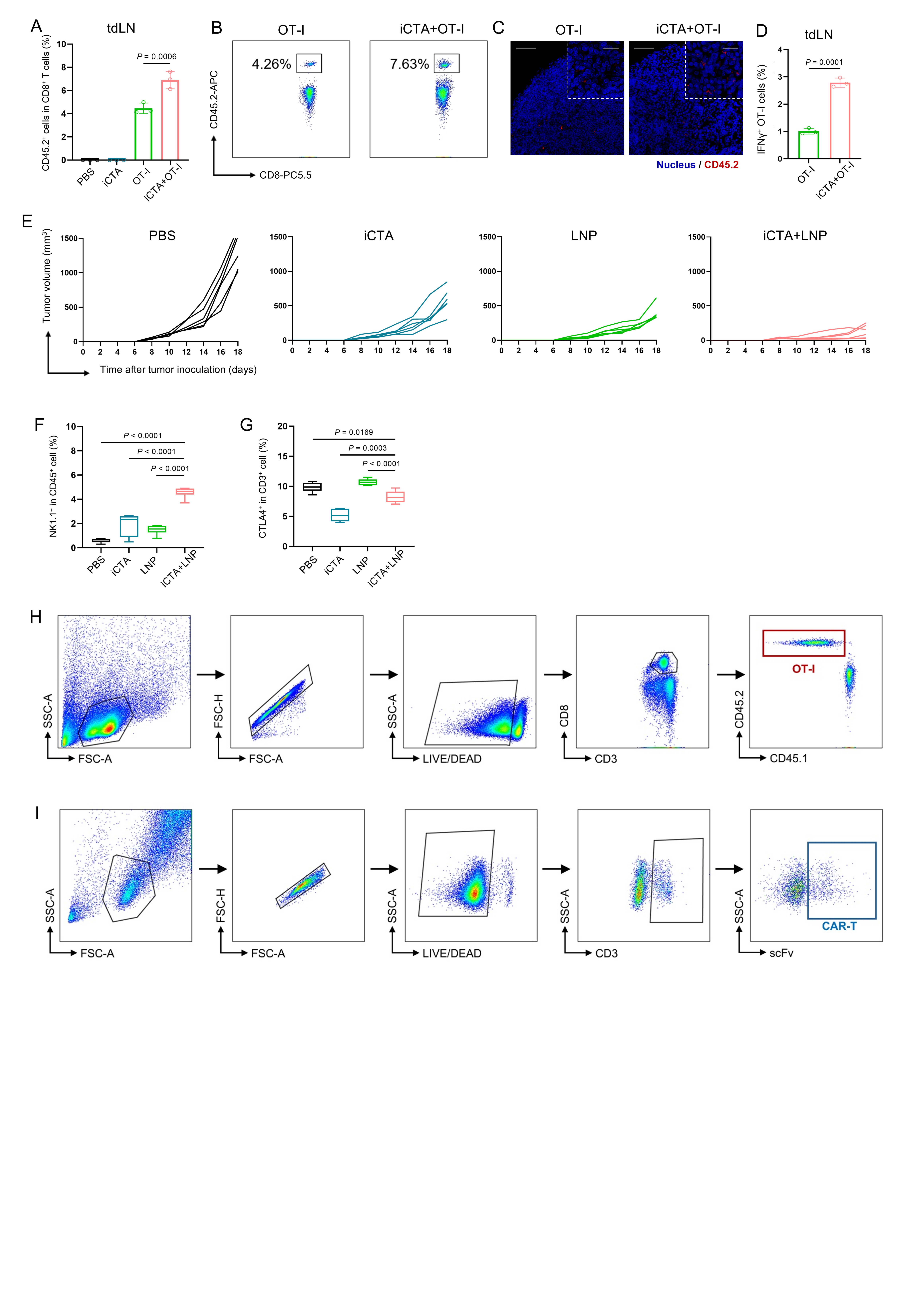

**Fig S4. Evaluation of the therapeutic efficacy of combination therapy**

(A, B) Flow cytometric evaluation of CD45.2⁺ OT-I cell in tdLNs. (A) Quantification of percentage among CD8⁺ T cells and (B) representative flow cytometry plots showing CD45.2⁺ OT-I cells in tdLN (n = 3).

(C) Representative immunofluorescence images showing the distribution of CD45.2⁺ OT-I cells (red) and cell nuclei (blue) in tdLNs. Left scale bars, 50 µm; right scale bars, 20 µm.

(D) Flow cytometric analysis of intracellular IFNγ expression in CD45.2⁺ OT-I cells from the tdLN (n = 3).

(E) Individual tumor growth trajectories for each mouse from day 0 to day 18 after tumor challenge (n = 6).

(F) Flow cytometric evaluation of the proportions of NK cells among CD45⁺ cells in the TME (n = 6).

(G) Percentage of CTLA4⁺ cells in T cells (n = 6).

(H, I) Flow cytometry gating strategies for OT-I cells (H) and CAR-T cells (I).

Data in (A), (D), (F), and (G) are presented as the mean ± SD. Statistical significance was calculated via one-way ANOVA with a Tukey post-hoc test (A, F, and G) or two-tailed Student’s *t* test (D).

|  | HCT (%) | MCV (fL) | MCH (pg) | MCHC (g/L) | PLT (K/μL) |
| --- | --- | --- | --- | --- | --- |
| Al | 56.6±1.7 | 52.5±0.3 | 15.2±0.2 | 288.4±3.8 | 822.0±97.0 |
| Mn | 50.7±2.6 | 52.2±0.2 | 15.5±0.3 | 295.8±5.9 | 787.2±48.5 |
| iCTA | 53.5±3.3 | 51.6±0.6 | 15.2±0.4 | 294.8±6.4 | 748.6±68.7 |
| PBS | 50.7±1.1 | 52.3±0.3 | 15.5±0.2 | 296.0±3.3 | 786.2±57.8 |

**Table S1. Hematological parameters**
